# CRISPR-Cas9 targeted disruption of the *yellow* ortholog in the housefly identifies the *brown body* locus

**DOI:** 10.1101/105023

**Authors:** Svenia D. Heinze, Tea Kohlbrenner, Domenica Ippolito, Angela Meccariello, Alexa Burger, Christian Mosimann, Giuseppe Saccone, Daniel Bopp

## Abstract

The classic *brown body* (*bwb*) mutation in the housefly *Musca domestica* impairs normal melanization of the adult cuticle. In *Drosophila melanogaster*, a reminiscent pigmentation defect results from mutations in the *yellow* gene encoding dopachrome conversion enzyme (DCE). Here, we demonstrate that the *bwb* locus structurally and functionally represents the *yellow* ortholog of *Musca domestica*, *MdY*. In *bwb* Musca strains, we identified two mutant *MdY* alleles that contain lesions predicted to result in premature truncation of the MdY open reading frame. We targeted wildtype *MdY* by CRISPR-Cas9 RNPs and generated new mutant alleles that fail to complement existing *MdY* alleles, genetically confirming that *MdY* is the *bwb* locus. We further found evidence for Cas9-mediated interchromosomal recombination between wildtype and mutant *bwb* alleles. Our work resolves the molecular identity of the classic *bwb* mutation in *Musca domestica* and establishes the feasibility of Cas9-mediated genome editing in the Musca model.

## Introduction

The so-far unidentified *brown body* (*bwb*) locus in the housefly *Musca domestica* was named after a recessive loss-of-function phenotype in which the adult cuticle manifests in a brown color rather than the wildtype black pigmentation **(Fig. 1a).** The absence of black coloration in mutant *Musca* has been proposed to result from impaired synthesis and incorporation of the black pigment melanin during pupal stages [1]. In insects, melanization of the cuticle is widely common and contributes to the diverse coloration patterns that are the most visible features of the outer morphology. Most insights into the pathway that produces and incorporates melanin into the insect cuticle comes from studies in *Drosophila melanogaster* [2] [3] [4]. The melanin pathway starts with conversion of tyrosine to Dihydroxyphenylalanine (DOPA) by *tyrosine hydroxylase* (TH). DOPA in turn is converted to dopamine by *dopa carboxylase (DDC)*. Both substrates are used as precursors for production of black melanin. In *Drosophila*, the dopachrome conversion enzyme (DCE) catalyzes steps downstream of TH and DCC in melanin production; however, whether DCE is involved in only in DOPA, only in dopamine conversion, or in both, remains unclear. Loss of function in the *yellow* gene (*y*) that encodes *Drosophila* DCE causes a lack of melanin incorporation and results in a yellowish overall appearance of the cuticle. Similar phenotypes have been observed in other insects such as the lepidopteran *Bombyx mori, Papilio xuthus,* and the coleopteran *Tribolium castaneum* [5] [6] [7] [8] [9]. Loss of activity of the corresponding *yellow* orthologs in these species drastically lowers the synthesis of melanin, causing regions of the body that are normally black to display a lighter coloration.

**Figure 1.**
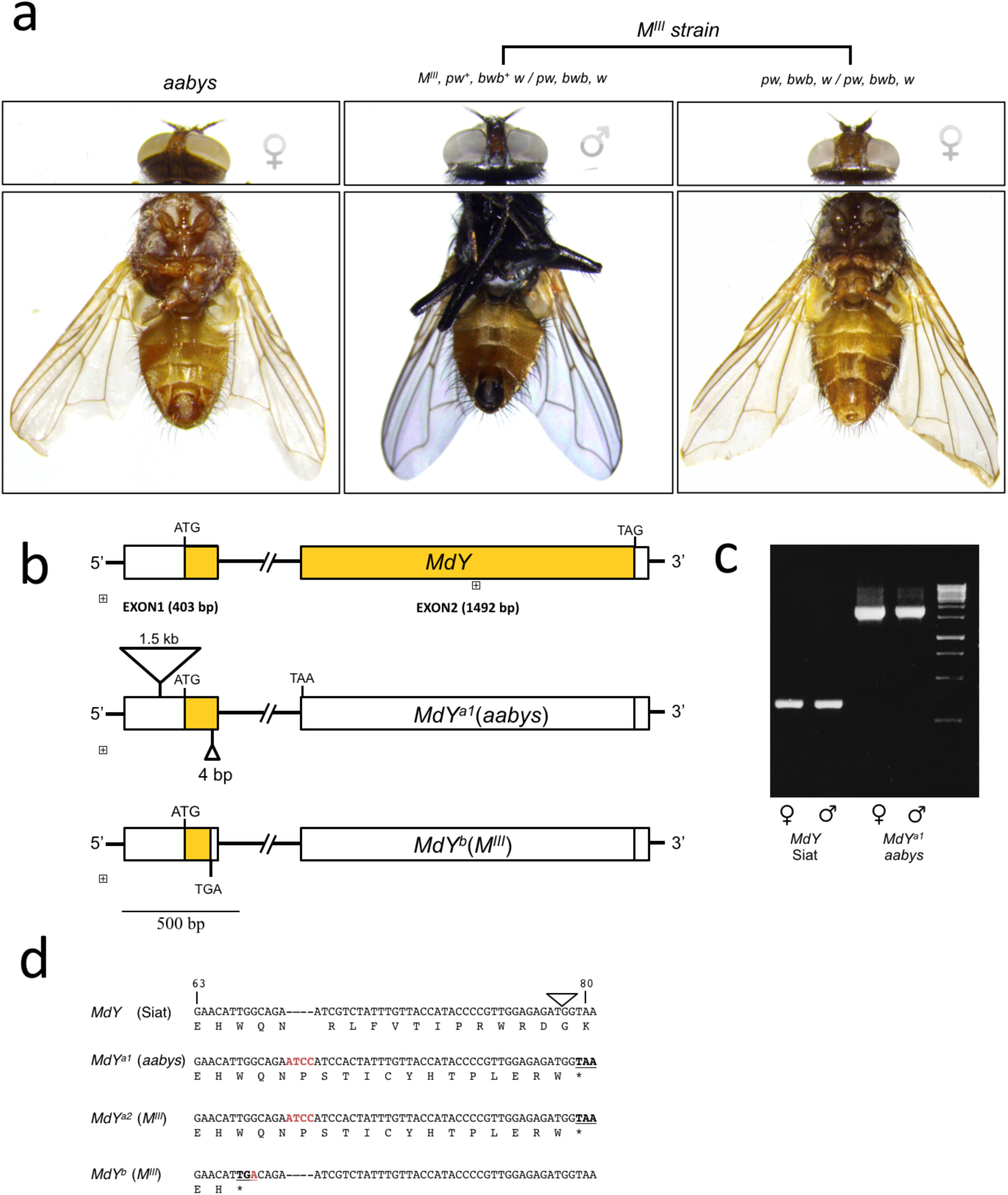
*bwb* phenotype is associated with nonsense mutations in MdY. **a)**Phenotypes are displayed from left to right: multimarked *aabys* fly, *bwb* mutant female from *M*^*III*^strain and *bwb* wildtype male from *M*^*III*^ strain. Genotypes are indicated; *bwb*: brown body, *pw*: pointed wings (notches along the edge of the wing, see arrow), *w*: white eyes.**b)**schematicdrawing of the *MdY* locus in the *bwb* mutant *aabys* strain and the Siat *bwb* wildtype strain. The aabys allele, *MdY*^*a1*^, contains a 1.5 kb insertion in the 5’ UTR and an additional 4 bp insertion in the ORF of exon 1. This frame-shift leads to a premature TAA stop in the 5’ end of exon 2. The Siat *MdY* allele has an intact ORF (boxed yellow). **c)** Genomic amplification with flanking primers Y-GAP1-F1 and Y-EXON1-R show that the 1.5 kb insertion is present in males and females of the *bwb* mutant aabys strain, but not in the *bwb* wildtype Siat flies. **d)** Affected part of the coding region (position 63 to 80) of the nonsense alleles of *MdY*. Deviations from the wildtype sequence (Siat) are marked in red and translational stops in bold. Location of intron is indicated with a triangle

In recessive *bwb*-mutant houseflies, the loss of normal black coloration affects all body parts. This phenotype closely resembles the phenotype observed in *yellow*-mutant *Drosophila*. As no other gene in the network of melanin genes is known to manifest this phenotype, we sought to investigate whether the *bwb* gene in the housefly is the structural and functional homolog of the *yellow* (*DCE*) gene in *Drosophila*. In addition to the correspondence in phenotypes, the *bwb* locus has been mapped to Chromosome III in Musca which corresponds to Muller element A, the X chromosome in Drosophila that harbors the *y* locus [10,11].

Here, we identified the locus mutated in *bwb* as the *Musca domestica* ortholog of the *Drosophila* gene *yellow* and refer to the gene as *MdY*. The *MdY* gene shows a high degree of similarity at the level of both protein sequence and gene structure. We find three mutant alleles of the *MdY* gene in *bwb*-mutant flies: the alleles *MdY*^*a2*^ and *MdY*^*b*^ feature sequence disruptions of the coding sequence, while a third allele *MdY*^*a1*^ is a compound allele featuring the coding lesion of *MdY*^*a2*^ and an additional 1.5 kb sequence insertion in the *5’ UTR*. Using CRISPR-Cas9, we generated a series of *de novo* loss-of-function alleles of *MdY* that all fail to complement original *bwb* alleles; these experiments mark, to our knowledge, the first report of the successful application of Cas9-based mutagenesis in *Musca domestica*. Altogether, we conclude that the *bwb* phenotype in Musca is caused by a lack of DCE activity normally provided by the *MdY* gene. Our findings clarify the molecular lesions in the classic *bwb* mutation and further underline the notion that *yellow* plays a conserved role in the melanin production pathway in dipteran species.

## Results

### 1. Characterization of the yellow ortholog in Musca domestica

Based on phenotype resemblance and mapping position, we hypothesized that *bwb* affects the so-far undescribed DCE ortholog in *Musca domestica*. To identify sequences homologous to the *Drosophila yellow* gene in Musca, we performed BLAST searches against the recently published genome of the multi-marked *aabys* strain that shows the *bwb* phenotype [12]. We recovered an annotated mRNA sequence (NCBI RefSeq *XM_011292650.1*) with a high degree of sequence similarity to *Drosophila yellow* (Supplementary data Fig. 1). Annotation of the *aabys* genome called this gene a pseudogene based on the lack of an intact open reading frame. Indeed, we detected a frame-shift starting 67 codons downstream of the first AUG start codon (**Fig. 1b**) [12]. Hence, the molecular nature of this allele already suggested that the *bwb*-mutant *aabys* strain carries a non-functional *yellow* variant. The mutated putative Musca *yellow* gene, which we named *MdY*, is located on *Scaffold18750* (502 kb) and is composed of two exons separated by a 35.6 kb-spanning intron (**Fig. 1b**). A *Musca* homolog of the *acheate* (*ac*) gene is present on the same scaffold separated by 156 kb, revealing a relatively close linkage of *yellow* and *ac* orthologs in *Musca.* This coupling is conserved in *Drosophila melanogaster* and sibling species, and previous work proposed that evolutionary variation of the *y-ac* region is reduced due to the selective fixation of one or more advantageous mutations in this region [13]. We next isolated *MdY* sequences from a wildtype strain *Siat* that displays a normal melanization pattern; the *MdY* sequence in *Siat* has an intact ORF of 522 amino acids, markedly lacking the 4 bp insertion that causes a frame-shift in the *aabys*-derived allele (**Fig. 1b and d**). The predicted MdY protein shares a high level of similarity with YELLOW proteins of other dipterans (Supplementary Figure 2a): as expected, *MdY* shares the highest level of identity (88%) with the predicted YELLOW protein in *Stomoxys calcitrans,* a close relative of the same family *Muscidae* (Supplementary data Fig. 2b).

To extend our analysis of mutant *bwb* alleles, we included the *M*^*III*^ strain that carries the male determining factor on Chromosome III tightly linked to wildtype alleles of *bwb* and *pointed wings* (*pw*), while females of this strain are homozygous mutant for *bwb* and *pw* (**Fig. 1a**). Analysis of *MdY* sequences isolated from *M*^*III*^ females unveiled that again the *bwb* allele that contains a 4 bp insertion at amino acid position 67. Unexpectedly, we also reproducibly detected a second mutant *MdY* allele in this strain, a nonsense mutation at amino acid position 65 (**Fig. 1d**). Therefore, the *M*^*III*^ laboratory strain carries two different loss-of-function alleles of *MdY*.

When mapping wildtype genomic *MdY* fragments against the corresponding *aabys* genomic sequences that harbor the 4 bp insertion in the coding sequence, we additionally identified a structural difference in the *5’ UTR* region upstream of the putative start codon (**Fig 1b**). Sequence analysis revealed the presence of a 1.5 kb insertion in the *aabys-*derived allele that is absent in the wildtype *Siat* strain (**Fig 1b and c**). A BLAST search against the Musca genome shows that this *aabys*-specific insertion shares 77% of sequence identity to an incomplete gene complement of the nicotinic acetylcholine receptor subunit-encoding *Mdalpha2* [14]. This 5’ UTR insertion in the mutant *MdY* allele of the *aabys* strain consequently represents a third, compound mutant allele of *MdY*.

Altogether, we identified two distinct coding-frame alleles in *bwb*-mutant *Musca domestica* strains, and a third compound lesion that introduced a 1.5kb insertion into the *5’ UTR* of one of the putatively inactive *MdY* loci. We refer to the original *aabys*-derived compound allele as *MdY*^*a1*^, and to the two *M*^*III*^-derived alleles as *MdY*^*a2*^ and *MdY*^*b*^ (**Fig. 1d, Supplementary data Fig. 3**). From our analysis of these mutants, we conclude that the *bwb* phenotype is associated with nonsense mutations in the *MdY* gene. These observations support the notion that the housefly DCE homolog is involved in the melanin production pathway.

### 2. CRISPR-Cas9-mediated disruption of MdY confirms causative association with bwb

We next performed targeted disruption of the wildtype *MdY* locus to confirm its predicted role in melanization of the housefly cuticle and to corroborate whether the *bwb* mutations are indeed loss-of-function alleles of *MdY*. To this end, we selected two target sites in the second exon of *MdY* for non-homologous end joining (NHEJ)-mediated disruption by the CRISPR-Cas9 system. Preassembled ribonucleoprotein complexes (RNPs) composed of purified Cas9 protein loaded with two different sgRNAs (*sgY2* and *sgY3*) were injected into early syncytial embryos. Both *sgY2* and *sgY3* target two different sites in coding exon 2 *separated by* 340 bp, (**Fig 2a**). As host strain, we used the *M*^*III*^ strain that carries the male determining factor on the chromosome III linked to wildtype alleles of *bwb* and *pw*. Females of this strain are homozygous mutant for both markers and are brown coloured with pointed-wings, while males are heterozygous and phenotypically wildtype for both markers ([15,16]) (**Fig. 2b**). This genetic background facilitates detection of possible somatic effects of *MdY* disruption in males that carry only one wildtype allele of *bwb* (**Fig. 1a**). The two sgRNAs were preloaded separately on purified recombinant Cas9 protein ([17], A.M. and G.S. unpublished results) and a 1:1 mix was micro-injected into 1 h old embryos of the M^III^ strain[18]. Of 2565 embryos injected with a 1:1 mix of solubilized Cas9 RNPs containing *sgY2*, and *sgY3*, we recovered 188 surviving adult houseflies. While 106 of these adults were males, none of the males displayed any patches of brown coloration, indicating that our targeting procedure does not introduce significant somatic mutation mosaicism. We proceeded with crossing these injected G0 males to *bwb*-mutant females of the same *M*^*III*^ strain (2 males and 6 females per cross) to screen for possible germline effects. Screening the F1 generation, we observed 17% of crosses that produced brown-colored *M*^*III*^ *pw*^*+*^ males that can only arise from paternal transmission of a mutant *bwb* allele (**Tab. 1, Fig. 2C**). This observation reveals that our Cas9 RNP-mediated mutagenesis protocol introduced *bwb* mutations by targeting the *MdY* gene in germ cells of the syncytial embryo. Of note, the injected males vary greatly in the proportion of potentially *MdY*-disrupted F1 individuals they sire (**Tab. 1),** revealing variable germline mosaicism resulting from Cas9 mutagenesis akin to observations in other model organisms [18].

**Figure 2.**
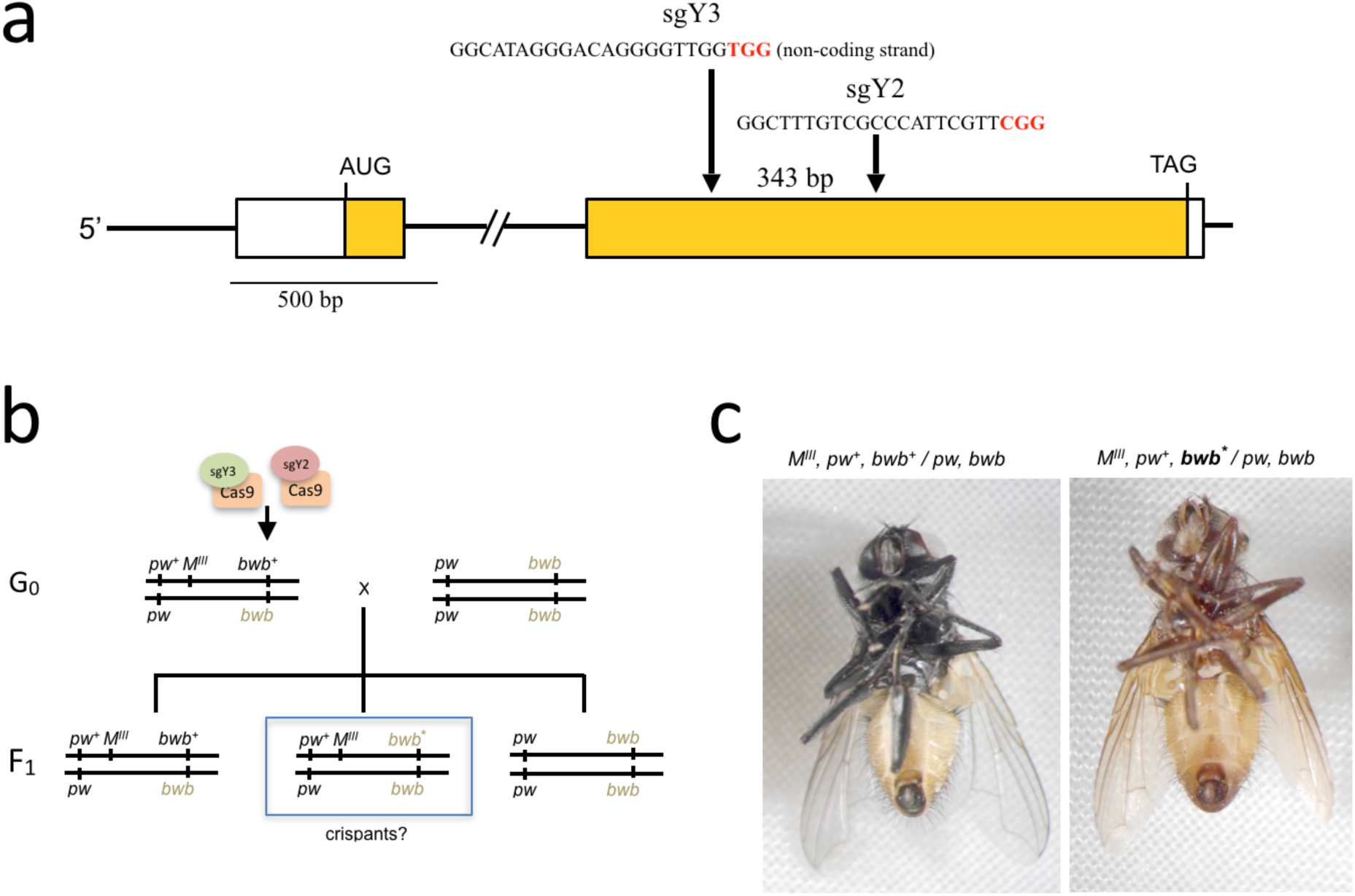
Strategy for CRISPR/Cas9 mediated disruption of *MdY*. **a)** A schematic of the *MdY* gene showing the positions of the two target sites in exon 2. Sequences used for the design of the two sgRNAs, sgY3 and sgY2, are indicated. Both sequences are flanked by a PAM motif (in red) and separated by 343 bp. **b)** Crossing scheme for screening mutational events affecting melanization. Injected G_0_ males (with a mix of sgY3-CAS9 and sgY2-CAS9) are crossed with *bwb* females and F1 is examined for occurrence of *bwb* males. **c)** Left a *bwb* mutant F1 male from line MdY#16 which is heterozygous for aCRIPSR induced 10 bp deletion in sgY3 over *MdY*^*b*^. Right an unaffected *bwb* wildtype F1 male from the same line with the paternal genotype (*MdY*^*+*^ over *MdY*^*b*^)

**Table 1.** Lines with bwb males in F1 generation. From a total of 43 crosses with 2 injected G_0_ males each, 14 lines with *bwb* mutant F1 males were recovered. MdY#29 and MdY#40 also produced *bwb*^*+*^ wildtype females. The numbers of F1 flies with different phenotypes are shown for each line. Presence of *M*^*III*^ indicates a male phenotype.

| line | $M^{III} pw^+ bwb^+$ | $pw bwb$ | $M^{III} pw^+ bwb$ | $pw bwb^+$ |
| --- | --- | --- | --- | --- |
| MdY#2 | 198 | 484 | 18 | 0 |
| MdY#4 | 212 | 183 | 1 | 0 |
| MdY#7 | 136 | 150 | 1 | 0 |
| MdY#9 | 152 | 136 | 3 | 0 |
| MdY#10 | 28 | 86 | 151 | 0 |
| MdY#13 | 110 | 124 | 12 | 0 |
| MdY#14 | 139 | 55 | 1 | 0 |
| MdY#16 | 179 | 174 | 21 | 0 |
| MdY#19 | 147 | 115 | 13 | 0 |
| MdY#29 | 142 | 136 | 68 | 10 |
| MdY#33 | 257 | 211 | 7 | 0 |
| MdY#36 | 163 | 138 | 4 | 0 |
| MdY#38 | 194 | 171 | 5 | 0 |
| MdY#40 | 81 | 62 | 2 | 1 |

To characterize the putative lesions induced in the *MdY* gene by sgRNAs *sgY2* and *sgY3*, we PCR-isolated genomic sequences of exon 2 from brown-bodied *M*^*III*^ *pw*^*+*^ F1 males by PCR and examined the targeted sites in sub-cloned and Sanger-sequenced fragments using CrispRVariants [19]. Overall, the position and extent of the induced lesions vary between individual lines but show a clear preference for lesions at the *sgY3* target site (**Table 2, Supplementary Fig. 4**). In seven lines (*MdY#2, MdY#9, MdY#10, MdY#13, MdY#16, MdY#33, MdY#36,* and *MdY#38*) we found small indels exclusively in the target site of *sgY3* (Table 2). In contrast, only line *MdY#14* carries an allele of *MdY* with a lesion (2 bp deletion) in the *sgY2* site (**Table 2, Supplementary Fig. 4**). In line *MdY#19*, we found evidence that both sites were targeted (Supplementary Fig. 4). The absence of a deletion spanning the two sites suggests that two dsDNA break events in our recovered alleles must have occurred at different times or with different kinetics, allowing the cellular repair system to independently join the breaks. In contrast, line *MdY#40* carries a large deletion of 1038 bp which removes both target sites and sequences downstream of *sgY2*; *MdY#40* likely had both sites simultaneously targeted by Cas9 and repaired to result in a larger deletion. Taken together, these data establish functional CRISPR-Cas9-mediated mutagenesis targeting the *MdY* locus in *Musca domestica* using *in vitro*-assembled Cas9-sgRNA RNPs.

**Table 2.** Overview of *MdY* lesions in *bwb* males. 14 CRISPR lines are listed with lesions detected in the target sites, ∆bwb sg3 and ∆bwb sg2. The majority of mutations caused by NHEJ occurred in the ∆bwb sg3 region ranging in size from 1 bp insertion to 174 bp deletions. In line MdY#38 sequence changes in and downstream of ∆bwb sg3 but we were unable to determine the extent of this putative lesion (nd). In lines MdY#29 and MdY#40 none of the isolated sub-clones harbored a visible lesion in the region of the two target sites. Since both lines also produced recombinant *pw, bwb*^*+*^ females, it is likely that the *pw*^*+*^*, bwb* males are products from a recombinant event rather than from a NJEH induced lesion.

| line | $\Delta bwb\ sg3$ | $\Delta bwb\ sg2$ |
| --- | --- | --- |
| MdY#2 | 1 bp insertion | - |
| MdY#9 | 174 bp | - |
| MdY#10 | 15 bp | - |
| MdY#13 | 11 bp | - |
| MdY#14 | - | 2 bp |
| MdY#16 | 10 bp | - |
| MdY#19 | 14 bp | 11 bp |
| MdY#29 | - | - |
| MdY#33 | 12 bp | - |
| MdY#36 | 43 bp | - |
| MdY#38 | extent not defined |  |
| MdY#40 | 1038 bp |  |

To test for additional complementation, we crossed brown males of lines *MdY#10* which carries a 15 bp deletion in the *sgY3* site of *MdY* and *MdY#16* containing a frame-shifting 10 bp deletion at the same site with brown females of the *M*^*III*^ strain carrying the *MdY*^*a2*^ and *MdY*^*b*^ alleles and the *aabys* strain, homozygous for the *MdY*^*a1*^ allele. In all crosses, all progeny displayed *bwb* phenotype. Lack of melanization in these animals is consistent with our hypothesis that *MdY* is required for proper pigmentation of the cuticle. Furthermore, non-complementation of the Cas9-induced *MdY* alleles with all mutant *bwb* alleles of the *aabys* and *M*^*III*^ strains confirms our initial hypothesis that *bwb* corresponds to the *yellow* ortholog of *Musca domestica.*

Two lines, *MdY#29* and *MdY#40*, were of particular interest, as we did not detect any sequence modifications at the two target sites in exon 2 (**Table 2**). Nonetheless, both these lines produced both brown *M*^*III*^ *pw*^*+*^ males and also *pw* females with normal melanization. The reciprocity of these sex-specific phenotypes suggests Cas9-mediated double-strand breaks in *MdY* induced an intragenic recombination event between the *bwb*^+^ allele on the *M*^*III*^ chromosome and the mutant *bwb* allele on the corresponding homolog. This event may have created a recombinant *MdY* allele with abolished activity on the chromosome that contains the *M* factor. In line with this hypothesis, we found brown males in line *MdY#29* that are unaffected at the two Cas9 target sites, but homozygous for the *MdY*^*b*^ signature (translational stop at position 65) (**Fig. 3a**). These sequences are likely the products of a reciprocal recombination event that may also have resulted in a reconstitution of a wildtype *MdY* allele on the non-*M* chromosome in females. Indeed, we found that melanized females are heterozygous for the *MdY*^*b*^ signature in exon 1. Moreover, in the mutant *bwb* males, we detected heterozygosity for allele-specific polymorphisms just downstream of the target site *sgY3* that correspond to the paternal genotype (**Fig. 3a**). We interpret these observations as evidence for Cas9-mediated DNA double-strand breaks at, or upstream of, the *sgY3* site that induced an intragenic recombination event between the wildtype *MdY* allele and the *MdY*^*b*^ allele *in trans* in germ cells of Cas9 RNP-injected heterozygous males (**Fig. 3b**). This data indicates that Cas9-mediated mutagenesis in *Musca domestica* can lead to break-point-guided recombination in injected germ cells.

**Figure 3.**
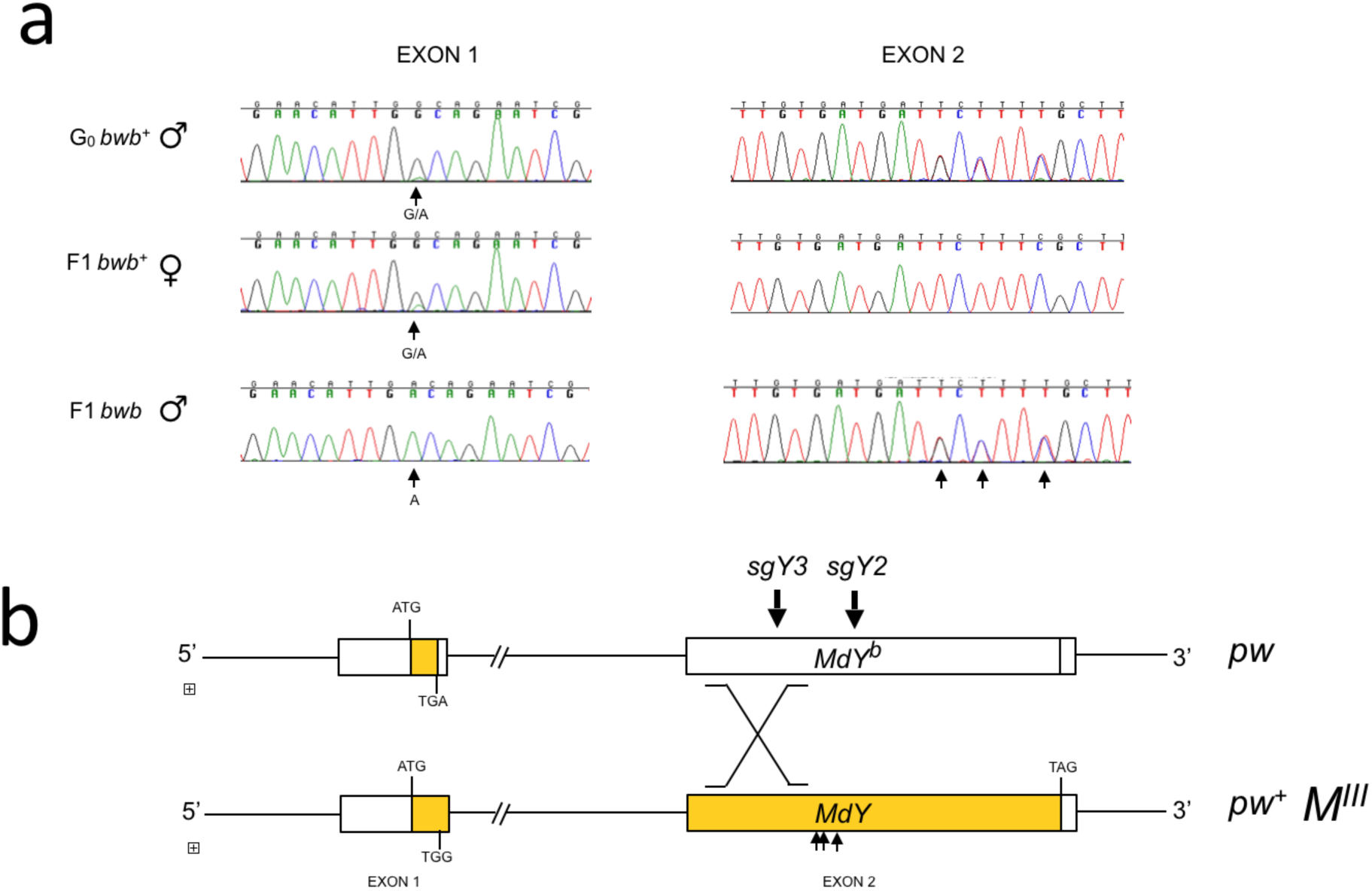
Intragenic recombination in *MdY* mediated by CRISPR/Cas9. **a)** Excerpts of chromatograms of exon 1 and 2 showing allele-specific polymorphisms. Mutant F1 *bwb* males are homozygous for the two variants in exon 1 (*MdY*^*b*^ genotype) amplified with primers Y-ORF-F3 and RE1, but heterozygous for the three variants in exon 2 like in the paternal wildtype *bwb* G_0_ male amplified with primers FE4 Y-ORF-R5. Arrows point to the allele-specific polymorphisms examined. **b)** This sequence analysis suggests that an intragenic recombination occurred downstream of the *MdY*^*b*^ specific TGA stop codon in exon 1 and upstream of polymorphisms examined in exon 2. Locations of the SNPs between the two target sites (sgY3 ad sgY2) are indicated with short arrows

## Discussion

Our work reveals that the classic *bwb* phenotype in houseflies is caused by mutations in the *Musca* homolog of the *DCE* gene, which encodes the enzyme that in *Drosophila* has been implicated in the process that converts DOPA and/or dopamine into black melanin. Functional studies in *Drosophila* have provided evidence that the DCE-encoding gene *yellow* is involved in global body pigmentation [20], whereas in the coleopteran *Tribolium castaneum* the loss of *yellow* only affects pigmentation of the hindwing [5]. Also, in the hemimetabolous *Oncopeltus fasciatus*, silencing of *yellow* affects only specific body parts such as abdomen and hindwings [21]. These observations led to the proposition that *yellow* belongs to a network of melanin synthesis genes which by differential deployment can generate a wide range of colors and spatial patterns [21]. Here, we present genetic evidence that, as in *Drosophila*, the *Musca* DCE homolog *MdY* is required for melanin production in the whole body. We base our conclusion of two major lines of evidence: First, complete absence of melanization found in two different *bwb* strains (*M*^*III*^ and *aabys*) correlates with homozygosity for predicted nonsense alleles of *MdY*. In these strains, we identified three mutant alleles *MdY*^*a1*^*, MdY*^*a2*^ and *MdY*^*b*^, all of which result in premature stop codons. We note that *MdY*^*a1*^ and *MdY*^*a2*^ only differ with regard to a 1.5 kb insertion which is present specifically in the *5’ UTR* of *MdY*^*a1*^. It is thus possible that these two alleles, both of which carry the same 4 bp insertion downstream of codon 67, have a common origin and that the 1.5 kb insertion has been acquired or lost later during a secondary mutational event. Second, we generated a set of new *MdY* alleles by adapting the CRISPR-Cas9 mutagenesis for disrupting the coding region in *MdY* exon 2. All of the new alleles have confined lesions at least in one of the two targeted sites. The majority of these mutations are deletions that remove parts of the coding sequence and generate frame-shifts. We hence consider these allelic variants to be enzymatically non-functional, if not null alleles of *MdY*. These Cas9-based *MdY* mutants fail to complement, and thus behave allelic to, the previously identified *MdY* variants found in *bwb*-mutant backgrounds.

The CRISPR-/Cas9 protocol using solubilized RNPs used in our study appears to be highly effective in the *Musca* germ line given that at least 1 of 6 RNP-injected males transmitted an *MdY* allele with a defined lesion at one of the targeted sites. Nonetheless, none of these F0 males displayed patches of non-melanized cuticle indicative of somatic mutation mosaicism. In *Drosophila*, *y* mutations behave cell-autonomously and even small mutant clones can be readily detected in the cuticle: when Cas9 mRNA was injected with sgRNA targeting *y* in *Drosophila* has strong somatic effects (86%) not only in hemizygous males with one wildtype target, but even in females with two wildtype copies of *y* [22]. This study pointed out that the efficiency of inducing somatic clones not only depends on the concentration of sgRNA injected, but more importantly on the selection of the site in the *y* gene that was targeted. It is thus conceivable that the sgRNAs used in our work were able to efficiently disrupt the gene in the germline cells but not in somatic cells. Germline transmission is a prerequisite for the investigation of any new mutation generated and, in this regard, our main objective was to recover fertile adults transmitting mutant *MdY* alleles. The lack of somatic effects in G_0_ adults can be a beneficial feature to avoid sterility that may be inflicted by the presence of mutant somatic tissue.

Recent work proposed that the CRISPR-Cas9 system can be used for genetic mapping by inducing targeted recombination events in meiotic and mitotic cells [23]. Our finding of an intragenic recombination event in line *MdY#29* suggests that Cas9 induced breaks allows recombination between homologs in the germ line of housefly males. This observation suggests a possible Cas9-mediated strategy for male meiotic mapping in future studies. In addition, homologous recombination (HR)-mediated repair of Cas9 induced double-strand breaks in *Musca* offers the opportunity to attempt template-based editing of genomic sequences for targeted knock-ins in the future.

To our knowledge, our study is the first report showing that Cas9 can be effectively deployed for NHEJ mediated-disruption of genes in *Musca domestica*, an important addition to the toolkit of molecular methods that have already been established in *Musca* for gene function analysis. Together with transient RNAi-based gene silencing [24,25] and stable germline transformation [16,26], this new genome editing system provides a means to investigate evolutionary diversification of developmental pathways such as the polymorphic sex determination system of the housefly [15,27]. Our successful attempt promises that this genome editing technology can be used in the housefly to study the function of any candidate gene of interest.

## Methods

### Rearing of houseflies

Rearing of larvae and adult flies has been described in [28,29]. Since low density of larvae on standard medium can cause substantial decrease in survival rates, we reared injected embryos and the surviving larvae on porcine manure. To dispose of mites and other parasites and to avoid contamination with eggs or larvae from wild-type populations, manure was stored at −70° C for at least two weeks prior to use.

### Strains of Musca domestica

1. wildtype strain was collected in Siat, Switzerland: females X/X*; bwb*^*+*^*/bwb*^*+*^ and males X/Y*; bwb*^*+*^*/bwb*^*+*^
2. multi-marked strain *aabys*: females X/X*; ac/ac; ar/ar; bwb/bwb; ye/ye; snp/snp* and males
3. autosomal *M*^*III*^ strain: females X/X*; pw, bwb, w/ pw, bwb, w* and males X/X*; pw*^*+*^, *M*^*III*^*, bwb*^*+*^*, w / pw, bwb, w* [16]

### Genomic DNA extraction

For genomic DNA extraction a single fly was collected in a 1.5 ml tube, frozen in liquid nitrogen and ground in 1 ml of extraction buffer (0.1 M Tris-HCl, pH 9; 0.1 M EDTA; 1% SDS and 1% of DMDC added freshly). After 30 min incubation at 70 °C, 140 µl of 8M potassium acetate was added and sample was gently inverted and incubated for 30 min on ice. After 15 min of centrifugation at 4°C at 13’000 rpm, supernatant was transferred to a new tube, and 550 µl of isopropanol was added. The mixture was centrifuged at RT for 5 min at 14’000 rpm and the supernatant was removed. The pellet was washed with 500 µl of 70% EtOH (-20 °C) and centrifuged at RT for 2 min at 14’000 rpm. The DNA pellet was finally dissolved in 30 or 50 µl of 10 mM Tris and 1 µl of RNaseA (10mg/ml) to remove RNA. Amplifications for sequence analysis were performed with following primers. For exon 1 we used forward Y-ORF-F3 (5’-TGCTGTGGACATTGGCAAGA -3’) and reverse RE1: (5’-TCTCATTCACATCCACACCGT-3’).

For exon2 we used forward FE4 (5’-CAGGTATACCAGCCACATTGA-3’) and reverse Y-ORF-R5 (5’-CTAATGATGGGCGGATGTGGA-3’). For insertion we used flanking primer forward Y-GAP1-F1 (5’-GGCCGAAGTGAGACAGAGAA-3’) and Y-EXON1-R (5’-CTAGTGGCGAAAAACCATTAA-3’).

### sgRNA synthesis and RNP complex assembly

SgRNA were designed using MultiTargeter Website (http://www.multicrispr.net). Possible OFF-target sites were excluded with the program Cas-OFFinder (http://www.rgenome.net/cas-offinder/) and by directly BLASTing selected sequences against the published housefly genome sequence [12]. Following templates for sgRNA production were generated:

Sgy2: 5’ -GAAATTAATACGACTCACTATA GGCTTTGTCGCCCATTCGTT GTTTTAGAGCTAGAAATAGC-3’

Sgy3: 5’ -GAAATTAATACGACTCACTATA GGCATAGGGACAGGGGTTGG GTTTTAGAGCTAGAAATAGC-3’

Common sgReverse (PAGE purified): 5’AAAAGCACCGACTCGGTGCCACTTTTTCAA GTTGATGGACTAGCCTTATTTTAACTTGCTATTTCTAGCTCTAAAAC AAC

For synthesis of sgRNA we followed the instructions of Megatranscript T7 kit (Ambion) using 400 ng of target template with a 5’ flanking T7 promoter as starting material. After RNA synthesis template was removed by incubating with TurboDNase (Mmessage Mmachine T7 Ultra Kit, Ambion) for 15 min at 37 °C [22].

Cas9 was expressed as an His-tagged protein and purified from bacteria (A.M. and G.S. unpublished results). The injection cocktail was prepared by mixing 1.5 µl purified Cas9 protein (9 mg/ml) with 2µl of sgRNA in 1.36 µl of 2M KCl in a total of 10µl. Prior injection the mix was incubated for 10 min at 37°C [18]. For injection we prepared a 1:1 mix of *sgY3*-preloaded Cas9 RNPs and *sgY2*-preloaded Cas9 RNPs.

### Microinjection of sgRNA-Cas9 complexes

Embryos of the *M*^*III*^ host strain were collected 1 hour after egg lay and chorion membrane was removed by incubating embryos in 3% sodium hypochlorite solution (NaOCl) for 1.5 min. Dechorionated embryos were then rinsed thoroughly with water and Ringer’s solution. Embryos were aligned on a cover slip with posterior ends pointing to injection site, dehydrated for 4 min in a silicagel chamber and then covered with 3S /10S (1:4) Voltalef oil (Prolabo). A glass needle was filled with the preloaded sgRNA-Cas9 mix which was injected into the posterior end of 0 to 1 hour old embryos. After injection excess Voltalef oil was carefully removed and cover slip were put on an agar plate overnight at 25°C. Surviving larvae were transferred after 24 hours to small beaker filled with porcine manure. G_0_ male individuals were collected shortly after eclosing and crossed with untreated virgin females of the *M*^*III*^ strain.

### Genomic analysis of CRISPR-Cas9 mediated lesions in MdY

To examine for the presence of lesions in *MdY* caused by NHEJ, genomic DNA of *bwb* mutant F1 males was extracted following the protocol described above. The region encompassing the two target sites was amplified with the following primer:

forward primers FE3: TCTGGCAAACCACAACAAGT or F4 CAGGTATACCAGCCACATTGA; reverse primers Y-ORF-R1: GACGAATGCCAACAACCCAC or Y-ORF-R5 CTAATGATGGGCGGATGTGGA

PCR products were purified with the Wizard^®^ Genomic DNA Purification Kit (Promega), subcloned in the pGEM®-T Easy Vector (Promega) and sent for Sanger sequencing (GATA BIOTECH).CrispRVariants and the generation of panel plots was performed from primary sequencing data as previously described. [18, 19]

## Acknowledgements

We thank Claudia Brunner for technical assistance, Raymond Grunder and Daniel Christen for help with the rearing of housefly cultures. We are grateful to Elena Chiavacci for her help with the target design and processing of CRISPR sequence data. This work was supported by the Canton of Zürich, a Schweizerischer Nationalfonds zur Förderung der Wissenschaftlichen Forschung (SNSF) professorship [PP00P3_139093] and a Marie Curie Career Integration Grant from the European Commission to C.M.; a University of Zurich URPP Translational Cancer Research Seed Grant to A.B. G.S. and A.M. were visiting researchers at University of Zurich (November 2015), supported by University Federico II of Naples (International Exchange Program to GS and by the Biology PhD program of University Federico II of Naples to AM).

## Authors contributions statement

DB. and GS conceived the experiments, SDH, TK, DI, AM. and AB. conducted the experiment(s). All of the authors discussed the data and helped manuscript preparation. DB, GS, and CM wrote the manuscript with intellectual input from all authors.

---

**Supplementary Figure 1.**
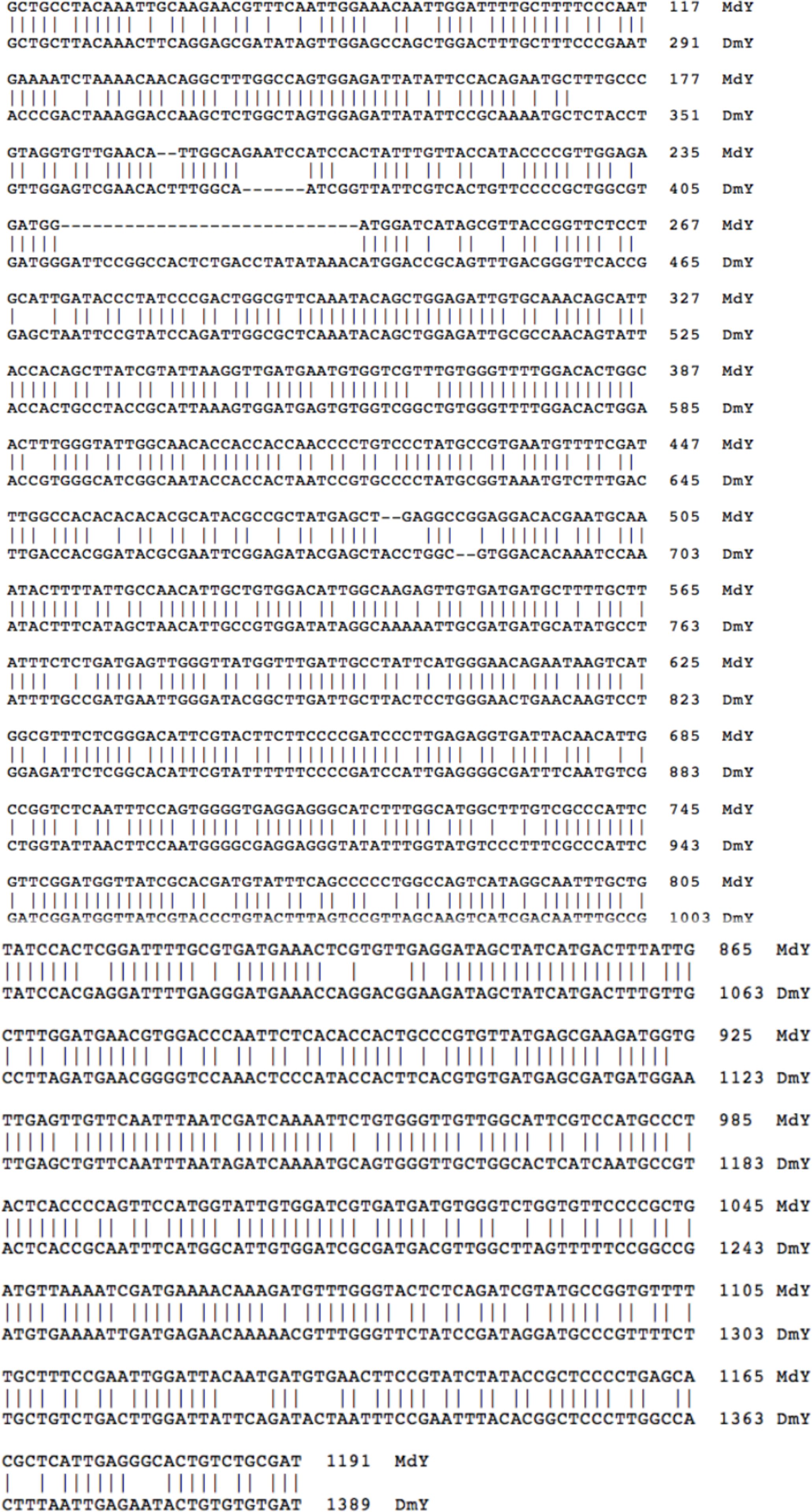
Nucleotide alignment of Drosophila *yellow (DmY)* and Musca *yellow* (MdY). The sequences used for alignment are XM_011297481.1 for Musca domestica yellow mRNA and NM_057444.3 for Drosophila melanogaster yellow mRNA. Overall identity is 74%

**Supplementary Figure 2.**
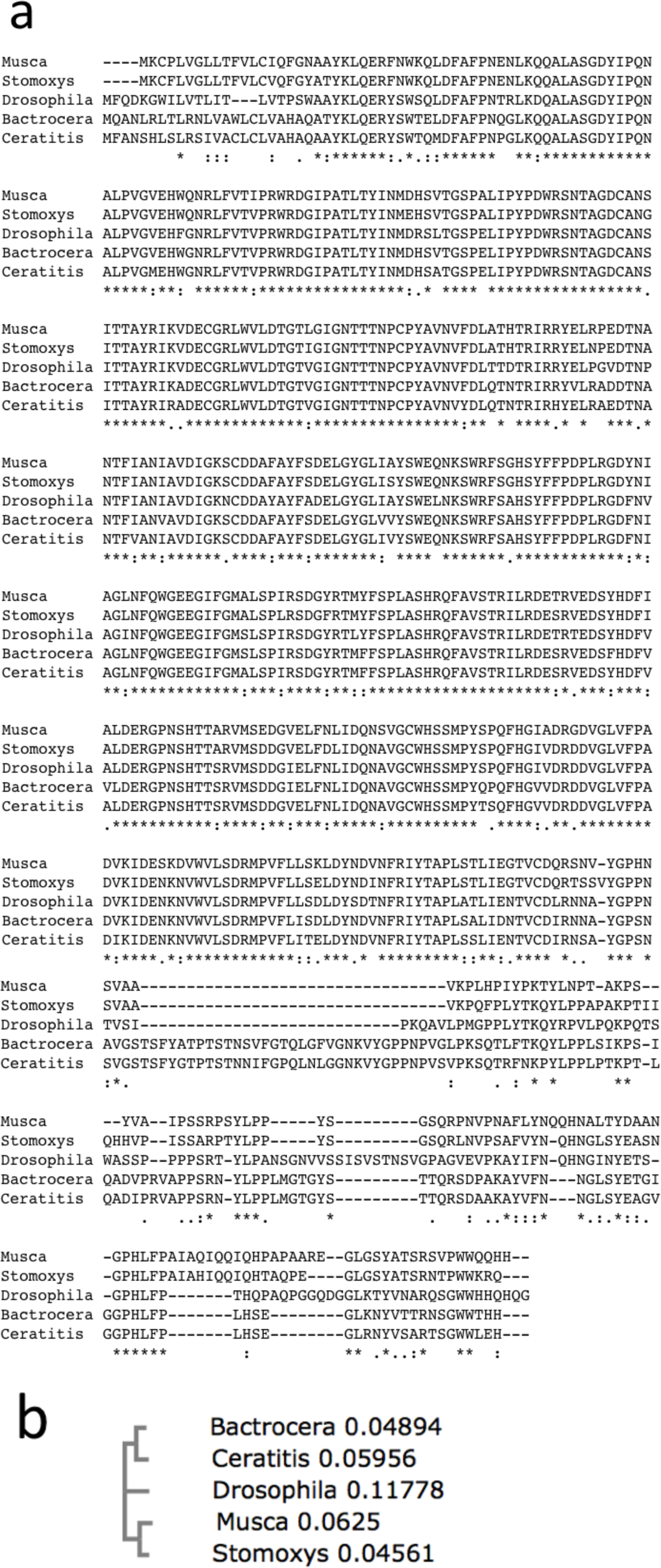
Amino acid sequence alignments of yellow orthologs found in higher dipterans. a) Clustal Omega alignment was performed with the following GenBank entries: Drosophila melanogaster yellow protein ID NP_476792.1, Stomoxys calcitrans yellow protein ID XP_013112765.1, Bactrocera oleae yellow protein ID XP_014099614.1, Ceratitis capitata yellow protein ID XP_004521097.1 b) The resulting Neighbour-joining tree with real branch length is depicted

**Supplementary Figure 3.**
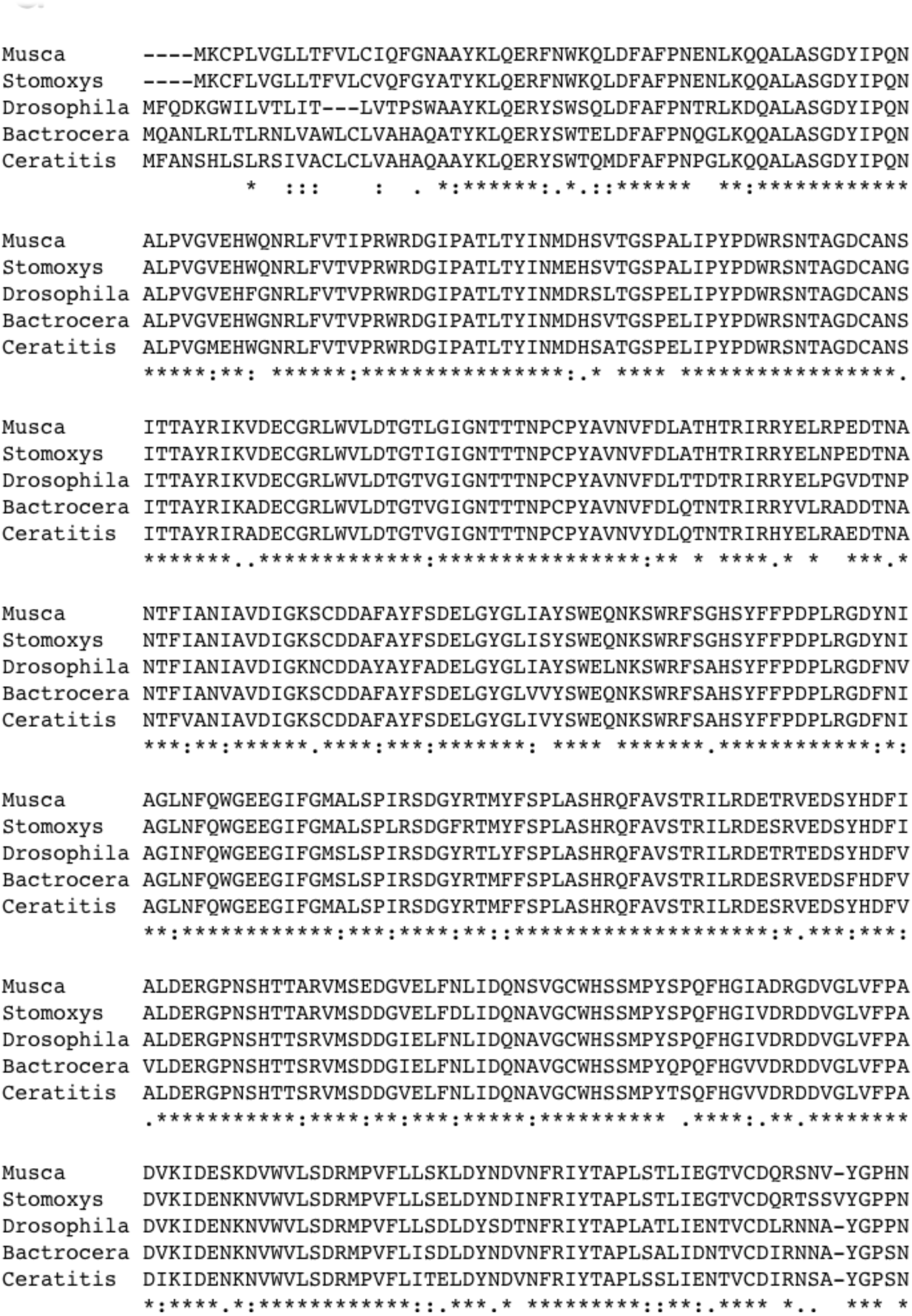
Nucleotide alignment of 2 different nonsense MdY alleles found in the *aabys* strain and in the MIII strain.

**Supplementary Figure 4.**
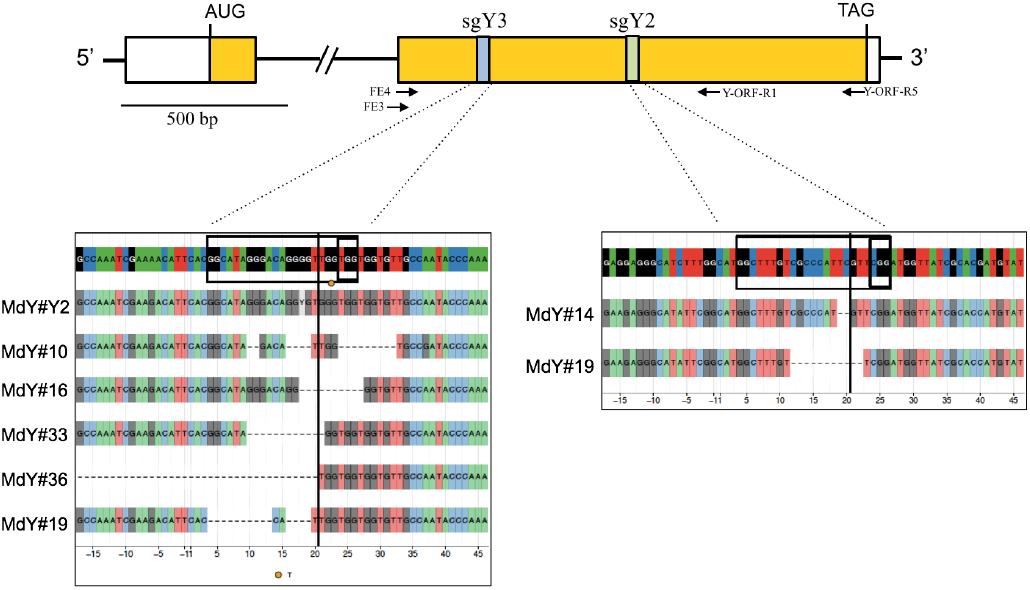
Compilation of lesions generated in targeted sites. Primers used for amplification and subcloning of fragments that were sequences are indicated with small arrows. CrispRVariants and the generation of panel plots was performed from primary sequencing data as described in [18, 19]

